# Neuromodulatory systems partially account for the topography of cortical networks of learning under uncertainty

**DOI:** 10.1101/2025.07.31.667879

**Authors:** Alice Hodapp, Florent Meyniel

## Abstract

Learning in dynamic and stochastic environments is notoriously difficult. Neuromodulatory systems may shape this process, thereby constraining where learning-related neural activity emerges according to the spatial distribution of receptors and transporters across the brain. However, the extent of this constraint, and which neuromodulatory systems contribute the most, remains unclear. Here, we focus on fMRI data from four probabilistic learning studies that was combined with a Bayesian ideal observer model. This model formalizes latent computational variables such as confidence and surprise that drive human learning. We show that the functional correlates of confidence, and to a lesser extent surprise, exhibit strong spatial invariance across tasks. This invariance suggests that these functional correlates of learning reflect a stable cortical organization largely independent of sensory modality and task structure. We found that this invariance aligns with the cortical chemoarchitecture using 20 PET-derived receptor and transporter density maps. Distinct receptor and transporter patterns mapped onto confidence- and surprise-related activity, including both known catecholamine pathways and novel opioid associations. Together, our findings provide evidence for a neuromodulatory account of adaptive learning and offer receptor-level hypotheses. More broadly, our approach provides a general framework to understand how neuromodulatory systems shape diverse cognitive processes.

## 1 Introduction

Learning in unstable and stochastic environments requires constantly deciding whether to update our expectations based on new observations. Deviations from what we expect may simply reflect random variability, or they may signal a meaningful change that warrants revising our estimates. These two situations call for very different learning responses. For example, when noticing that your train ride took longer than usual, you should update your estimated typical commute time if this observation denotes a hidden change, e.g. a new train schedule, and not update it if it simply arises from some expected variability, e.g. a routine signal failure. Bayesian inference provides an ideal and general solution to the updating problem: the amount of update should be guided by the amount of information the observation conveys about the latent parameter to estimate. In particular, observations that are more surprising (because they deviate more from the learned parameter) should be used to update more strongly, but this update should be smaller when the confidence about the parameter estimate is higher. Among other important factors, surprise and confidence therefore both regulate how much is learned from a single observation. Interestingly, both human behavior (1; 2) and neural data (e.g., 3; 4; 5; 6) show signatures of this Bayesian learning.

Previous studies indicate that on a neural level, neuromodulation regulates how much is learned from an observation. Specifically, major neuromodulatory systems such as dopamine, acetylcholine, norepinephrine and serotonin, have been proposed to implement the regulatory roles assigned to surprise and confidence in learning under uncertainty (7; 8; 9). These neuromodulatory systems project from brain stem nuclei to the basal ganglia, the cerebellum, and the cortex. They influence neurons and circuits by changing synaptic strength, excitability of neurons, and networks (e.g., 10; 11; 12).

The effect of neuromodulators greatly depends on the receptors onto which they bind, and on transporters. (13; 14; 15). Neurotransmitter receptors and transporters vary widely not only in terms of their affinity, timescales, and downstream effects on neuronal excitability, but also in terms of their topographies (their spatial distributions across the brain) (13; 15; 16). Motivated by prior research on the role of receptor and transporter function (14; 17; 18; 19), we here focus on the hypothesis that neuromodulation is a domain general mechanism of learning, and that therefore topographies of receptors and transporters across the brain should put constraints on the topography of neural activity during learning. More specifically, the cortex-wide activity patterns that reflect key computational variables of learning, namely surprise and confidence, should align to some extent with receptor and transporter density maps. This alignment is expected to be only partial, since other factors are known to provide additional constraints on the spatial organization of neural activity (e.g., connectivity, architecture, function; 20). Importantly, by enabling the identification of the receptors/transporters that best explain learning-related topographies, this approach also provides insight into the neuromodulatory systems that underlie learning. This is because the receptors that contribute most strongly to the observed spatial constraints likely point to the modulators involved in adaptive learning.

To investigate whether neurotransmitter receptors and transporters shape the topographical organization of domain-general learning effects, we analyzed fMRI data from multiple learning tasks. The tasks vary in terms of sensory modality (visual vs. auditory), generative structure (Bernoulli vs. Gaussian distribution vs. transition probabilities), and estimate to be learned (probability vs. reward magnitude). A prerequisite to our approach was to establish that the topographies of learning-related fMRI effects are indeed constrained. Therefore, in a first step, we demonstrated that cortical topographies of the effects of surprise and confidence were largely invariant across tasks. We then tested whether neuromodulatory systems could contribute to this invariance. To do so, we predicted the topography of functional MRI (fMRI) effects from receptor and transporter densities measured with positron emission tomography (PET). This analysis revealed that receptor/transporter densities partially account for the organization of the cortical networks engaged in learning under uncertainty. Finally, we identified specific receptors and transporters linked to surprise and confidence. While some associations correspond to previously established neuromodulatory pathways in learning, (7; 8), others point to novel relationships, underscoring the potential of receptor-level analyses to generate new hypotheses.

## 2 Results

To evaluate our hypothesis in a generalizable manner, we used a total of four different learning studies. Functional MRI data was recorded while participants performed a probability (Study 1-3, Figure 1A) or magnitude (Study 4, Figure 1B) learning task based on a sequence of events. Study 1 comprises 7T fMRI data (21) from participants presented with a visual sequence composed of two Gabor patches drawn from a Bernoulli process. Study 2 consists of yet unpublished 3T data in which participants were presented with an auditory sequence of two tones, determined by a Bernoulli process. Study 3 consists of 3T data (22) from participants presented with a sequence generated based on first-order transition probabilities. Auditory and visual sequences alternated in this study. Study 4 consists of yet unpublished 3T data from a magnitude (reward) learning task (23). Participants performed a sequence of choices in a two-armed bandit task trying to maximize cumulative reward, which required learning the underlying mean reward level of each option from Gaussian-distributed outcomes. In all studies, the latent generative parameter(s) changed occasionally and abruptly.

### 2.1 Bayesian inference accounts for subject behavior

Bayesian inference reportedly provides a good account of both probability learning (24) and magnitude learning (23; 25). Some internal variables of the Bayesian algorithm, such as confidence, are even accessible to introspection (4; 21; 22; 23; 26). Across the datasets analyzed here, participants were intermittently asked to report their confidence and, in most studies, their current probability or reward estimates during the course of learning. Consistent with previous findings, subjective confidence closely tracked the ideal observer in all four studies (Study 1: mean Pearson *r* = .16, *t*(25) = 5.9, *p* = 3.6× 10^*−*6^; Study 2: *r* = .18, *t*(54) = 8.8, *p* = 8.0 ×10^*−*12^; Study 3: *r* = .20, *t*(20) = 6.90, *p* = 1.1 ×10^*−*6^; Study 4: *r* = .29, *t*(55) = 14.60, *p* = 10^*−*19^), as visualized in Figure 1C. We also observed close correspondence between model- and subject-derived estimates of probability (Study 1: *r* = .79, *t*(25) = 31.8, *p* = 9.99 ×10^*−*22^; Study 2: *r* = .69, *t*(54) = 16.8, *p* = 1.0 ×10^*−*22^; Study 3: not available) and of reward magnitude (Study 4: fraction matching = .61; comparison with chance level (0.33): *t*(55) = 19.20, *p* = 10^*−*27^), visualized in Figure 1D. For an overview of linear regression results, see supplementary table 1. All subsequent analyses use trial-by-trial estimates of surprise and confidence computed by the ideal observer model. This is necessary since these quantities are not measured behaviorally on a trial-by-trial basis. The choice of an ideal observer model is motivated by first principles (2; 27), as well as the reasonable alignment between this model and the occasional reports of participants.

### 2.2 Confidence and surprise show invariant cortical organization

Turning to the main goals of this study, we first quantified the extent to which computational variables of learning exhibit invariant fMRI effects maps across studies. For each stimulus in the sequences, we derived key quantities of Bayesian inference from the ideal-observer model (see Materials and Methods). Confidence was formalized as the log precision of the posterior distribution, and surprise as the log improbability of the observation (28). These variables entered a general linear model (GLM) applied uniformly across studies (see Methods for covariates), and the resulting coefficients in each voxel of the brain constitute the subject-level effect maps used in all subsequent analyses.

Group-level z-statistic maps are shown in Figure 2. Confidence effects (Figure 2A) were highly consistent across studies, with strong overlap of negative clusters in the posterior intraparietal sulcus, precuneus, and anterior insula, and additional overlap in the middle frontal gyrus (Figure 2B). Surprise effects also exhibited cross-study overlap, with a core positive cluster in the right precentral sulcus/inferior frontal gyrus pars opercularis, as well as task-specific activations in sensory cortices and bilateral anterior insula in the reward-learning study (Figure 2D,E).

**Figure 1.**
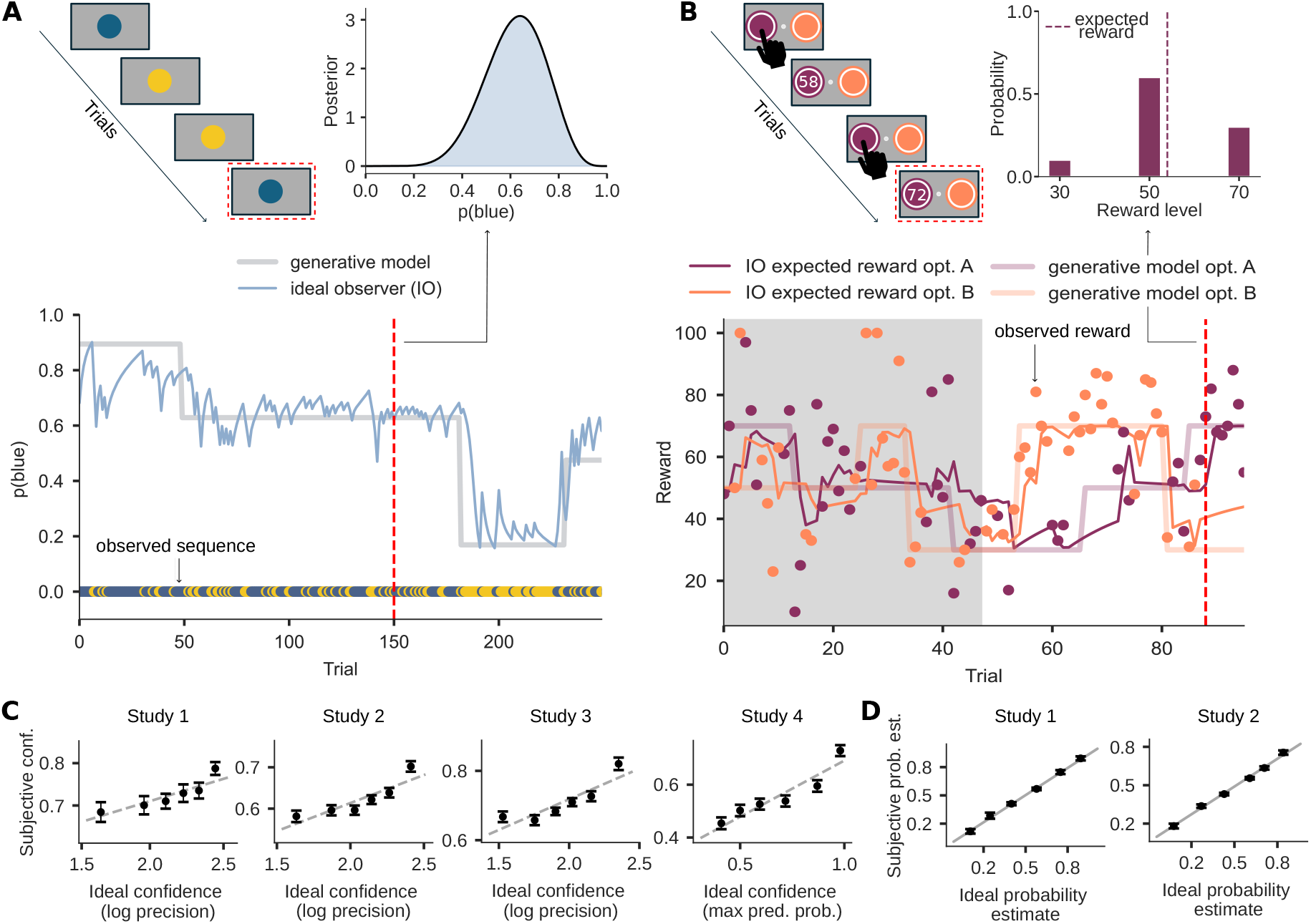
Tasks and behavior. A) Example probability learning task (similar to Study 1, 2). Top left: Four example trials. Bottom: Participants are presented with a sequence of two stimuli (here illustrated with blue and yellow dots). The generative probability p(blue) changed unpredictably (see gray line), the blue line represents the estimates of an ideal observer. Top right: Model posterior over p(blue) at the trial indicated by the red line. Note that the stimuli shown here are just for illustration, actual stimuli were geometric patterns or tones in different studies. B) Reward learning task (Study 4). Top left: Two example trials. At each trial subjects choose between two options and observe the outcome (reward) of the selected option. Bottom: The latent mean reward levels (light orange and purple) underlying the two options could change unpredictably between 3 reward levels (30, 50, 70). The observation (dark orange or purple dots) for the chosen option is drawn from a Gaussian distribution. The dark orange and purple lines represent the estimates of an ideal observer. Top right: Example posterior probability following an observation. C) All studies contained occasional reports about the latent parameters. Reported confidence levels sorted in 6 bins of ideal estimates. Dots and error bars indicate the mean and SEM. The dashed line is a linear fit. Note that for the probability learning studies (Studies 1-3) subjective confidence (lowest and highest response bins set to 0 and 1 arbitrarily) and ideal confidence (in log SD unit) are on different scales. D) Reported probability estimates sorted in 6 bins of ideal estimates. Dots and error bars indicate the mean and SEM. The solid line indicates the identity. Probability reports are not available for Study 3 and 4.

**Figure 2.**
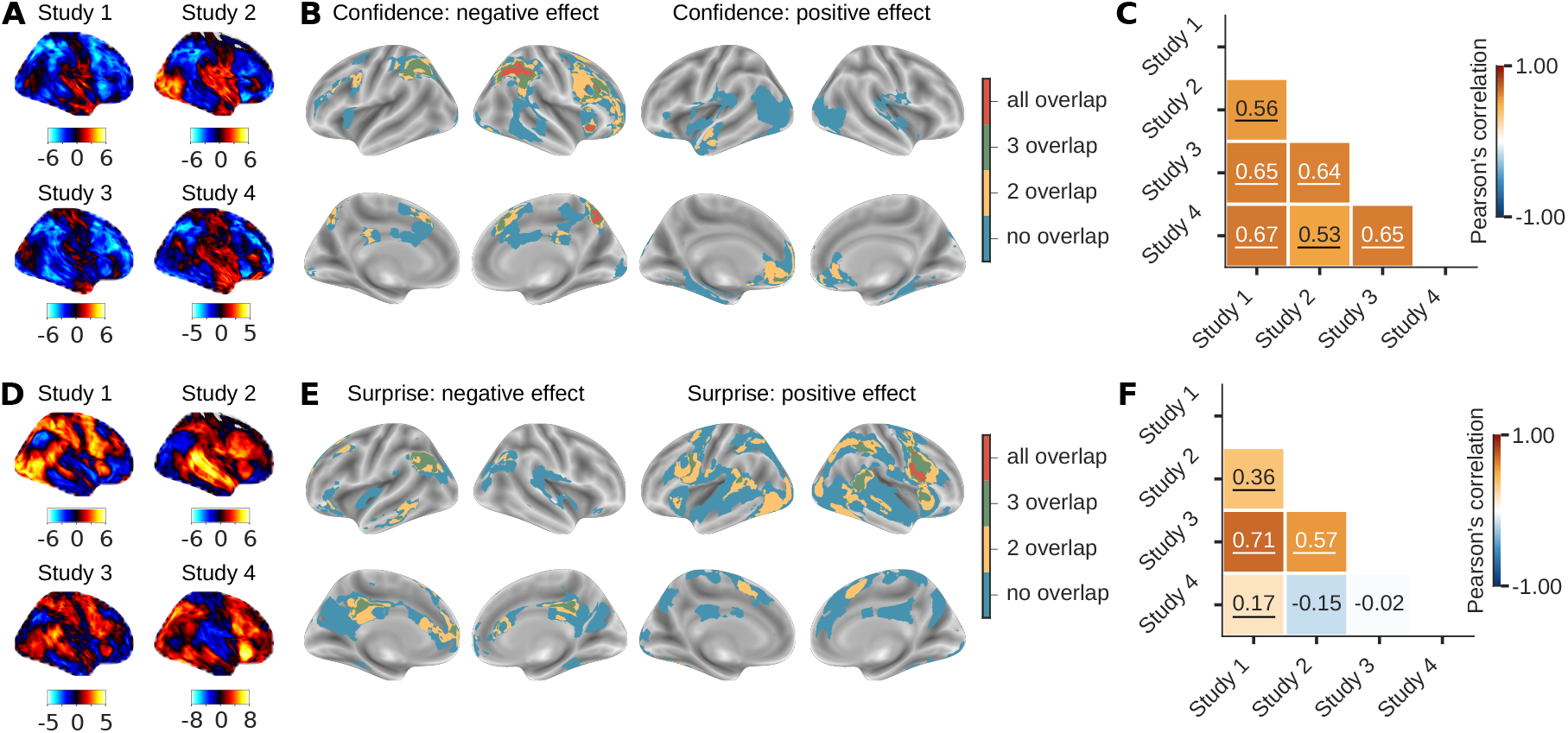
Effect maps are largely invariant across studies. A/D) Group-level z-statistic maps for confidence/surprise in the four studies. B/E) Brain regions where effects significantly correlated with confidence/surprise across studies. The colors correspond to the number of studies in which a region showed significant clusters (thresholded at *p*<0.001 and corrected for multiple comparisons across voxels at the cluster level with *p*_*FWE*_ <0.05). C/F) Correlations between effect maps across studies. Underlined correlation coefficients were significant in a comparison to a null model (all *p*= 0.001 at 1000 rotations)

To quantify invariance, we computed group-level correlations between effect maps for confidence and surprise across studies (Figure 2C,F). Confidence maps showed robust correlations across all tasks, significantly exceeding chance levels (all *p* = 0.001, FDR-corrected) as determined by spatial permutations that preserve the data distribution and spatial autocorrelation (‘spin test’; see Materials and Methods) (29; 30; 31). Surprise maps from the probability learning studies (Studies 1–3) were likewise positively correlated (all *p* = 0.001, FDR-corrected). In contrast, the surprise map from the reward-learning study (Study 4) did not correlate strongly with the others, consistent with differences in task design (see Discussion). Correlations between confidence and surprise maps are shown in the Supplement (Figure 5).

In sum, these analyses provide strong evidence for spatial invariance of confidence-related effects across all tasks, and for surprise-related effects within probability learning tasks. This invariance indicates that constraints exist on the topography of learning-related activity.

### 2.3 Neuromodulation partially accounts for spatial invariance of learning effects

We hypothesized that the observed invariance of confidence and surprise maps across tasks reflects neurobiological constraints imposed by neuromodulation (among others; 20). Specifically, we predicted that the topography of neurotransmitter receptor and transporter densities contributes to the topography of learning-related effects maps. We used receptor/transporter distributions across the cortex from published atlases, combining PET-derived whole brain density maps for 19 unique neurotransmitter receptors and transporters (14) with an additional *α*_2_ receptor density map (32).

To test our hypothesis, we predicted fMRI effect maps from receptor and transporter densities, using leave-one-subject-out cross-validation and multiple linear regression. The results show that the topography of receptors and transporters accounts for a significant proportion of variance in both confidence and surprise effect maps (Figure 3A). For each study–variable combination, the variance explained by the cross-validated models exceeded chance levels, as determined with the ‘spin test’ (see Materials and Methods): Confidence Study 1: *R*^2^= 0.035, *p*=0.017; Study 2: *R*^2^= 0.023; *p*=N.A; Study 3: *R*^2^= 0.036, *p*=0.0008; Study 4: *R*^2^= 0.009, *p*=0.044; Surprise Study 1: *R*^2^= 0.047, *p*=0.008; Study 2: *R*^2^= 0.036; *p*=N.A Study 3: *R*^2^= 0.022, *p*=0.038; Study 4: *R*^2^= 0.018, *p*=0.008 (FDR-corrected, one-sided; note that since Study 2 had incomplete cortical coverage, the ‘spin test’ is not applicable). Together, these results suggest that receptor topographies constrain the topography of learning-related activity.

**Figure 3.**
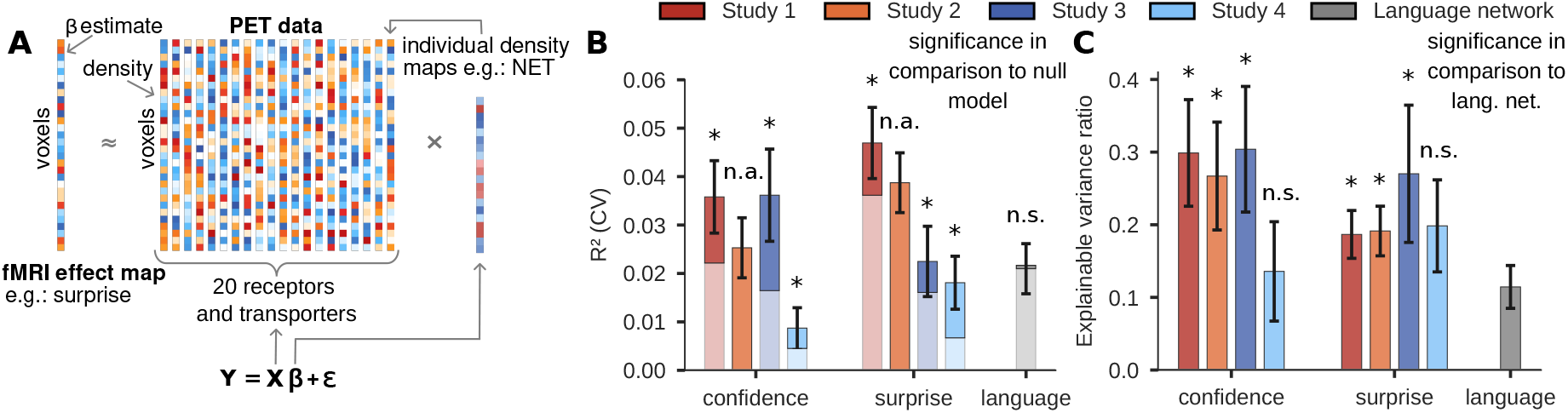
Learning-related effect maps can be predicted from receptor/transporter distributions. A) Analysis framework: Is there a linear relationship between a learning-related effect map and PET-derived density distributions of 20 receptors/transports in the cortex? B) Cross-validated *R*^2^ for a model including all 20 receptor/transporter distributions, by study and variable combination. The light part of each bar corresponds to the average performance of a spatial autocorrelation-preserving null model. The null model was not applicable to study 2, due to incomplete cortical coverage of the fMRI data. Error bars reflect the SEM. Asterisks denote significant models, compared to the null, in a paired t-test (FDR-corrected p <0.05, one-sided). n.a.: not applicable, n.s.: not significant. C) Mean cross-validated explained variance in ratio to the mean amount of variance that can be explained in a held out subject’s fMRI effect map by the other effect maps. Error bars reflect the bootstrapped SEM. Asterisks denote study/variable combinations in which the receptors/transporters explained a significantly larger fraction of variance than in the language network. Non-parametric permutation test on group level ratios (one-sided, FDR-corrected). n.s.: not significant.

We explored whether this relationship between effect maps and receptor distributions is specific to domains with strong neuromodulatory predictions, by applying the same analysis to a domain other than learning. To make this comparison meaningful, the alternative had to show robust, reproducible, whole-brain activation patterns across participants, yet lack strong neuromodulatory hypotheses. We used the fMRI language network (from a publicly available dataset: 33), as it met these criteria. Interestingly, receptor and transporter densities did not explain significantly more variance than the null model for the language network (Figure 3B). This finding is in line with the fact that neurobiological factors other than neuromodulation constrain the topography of the language network (34; 35; 36).

The metric used so far, cross-validated *R*^2^, is constrained by the amount of noise in the data, and thus difficult to interpret in absolute terms. We thus expressed mean model performance relative to the mean maximum explainable variance (Figure 3C; see Materials and Methods). For most learning-related study–variable combinations, receptor distributions explained a sizable proportion of variance (Confidence Study 1: 29.8 %; Study 2: 25.4 %; Study 3: 30.3 %; Study 4: 13.1 %; Surprise Study 1: 18.5 %; Study 2: 18.8 %; Study 3: 25.9 %; Study 4: 19.2 %; average across subjects). This proportion was lower for the language network (11.4 %), and significantly so compared to most other studies (Confidence Study 1: *p*=0.001; Study 2: *p*=0.001; Study 3: *p*= 0.001; Study 4: *p*= 0.68; Surprise Study 1: *p*=0.045; Study 2: *p*=0.023; Study 3: *p*=0.008; Study 4: *p*=0.07; FDR-corrected, one sided). These comparisons were tested for significance on group means with a non-parametric permutation test (see Material and Methods), the results for a test on the subject level (Man-Whitney U test) are reported in the the supplementary table 3 and lead to qualitatively similar conclusions. Importantly, these results do not suggest that receptor distributions are irrelevant for language processing. Rather, they support the hypothesis that neuromodulation plays a stronger role in shaping learning-related effects.

### 2.4 Specific receptors and transporters contribute to learning-related topography

Finally, we identified which individual receptors/transporters contributed most to the explained variance. To this end, we performed a dominance analysis (37) at the subject level (see Material and Methods). This analysis partitions the model fit among the predictors, while accounting for dependence or collinearity between them (Figure 6 shows pairwise correlations of topographies). Dominance reflects the relative contribution of predictors in an interpretable way. To facilitate comparison across models and studies, dominance values were expressed as proportions of the total model fit.

We first focused on confidence. Across all studies, dominance analysis revealed a consistent pattern of receptor/transporter contribution. Among all receptors, the topography of the *µ*-opioid receptor (MOR) contributed most strongly to explaining the topography of confidence effect maps (Figure 4A). Since dominance analysis ignores the sign of effects, we additionally inspected the regression weights in the regression model that uses all available predictors (Table 4). The MOR coefficient was significant across subjects in all probability learning studies (FDR-corrected p <0.05) and with a negative sign in each study. This indicates that regions with higher MOR density showed stronger activation under uncertainty (i.e., negative correlation with confidence), when taking into account all other receptors/transporters. Other receptors (5-HT_1B_, A_4_B_2_, CB_1_) also showed significant and consistent effects for each of the studies but contributed less to the model fit. Overall, receptor contributions were highly consistent across the probability learning studies.

**Figure 4.**
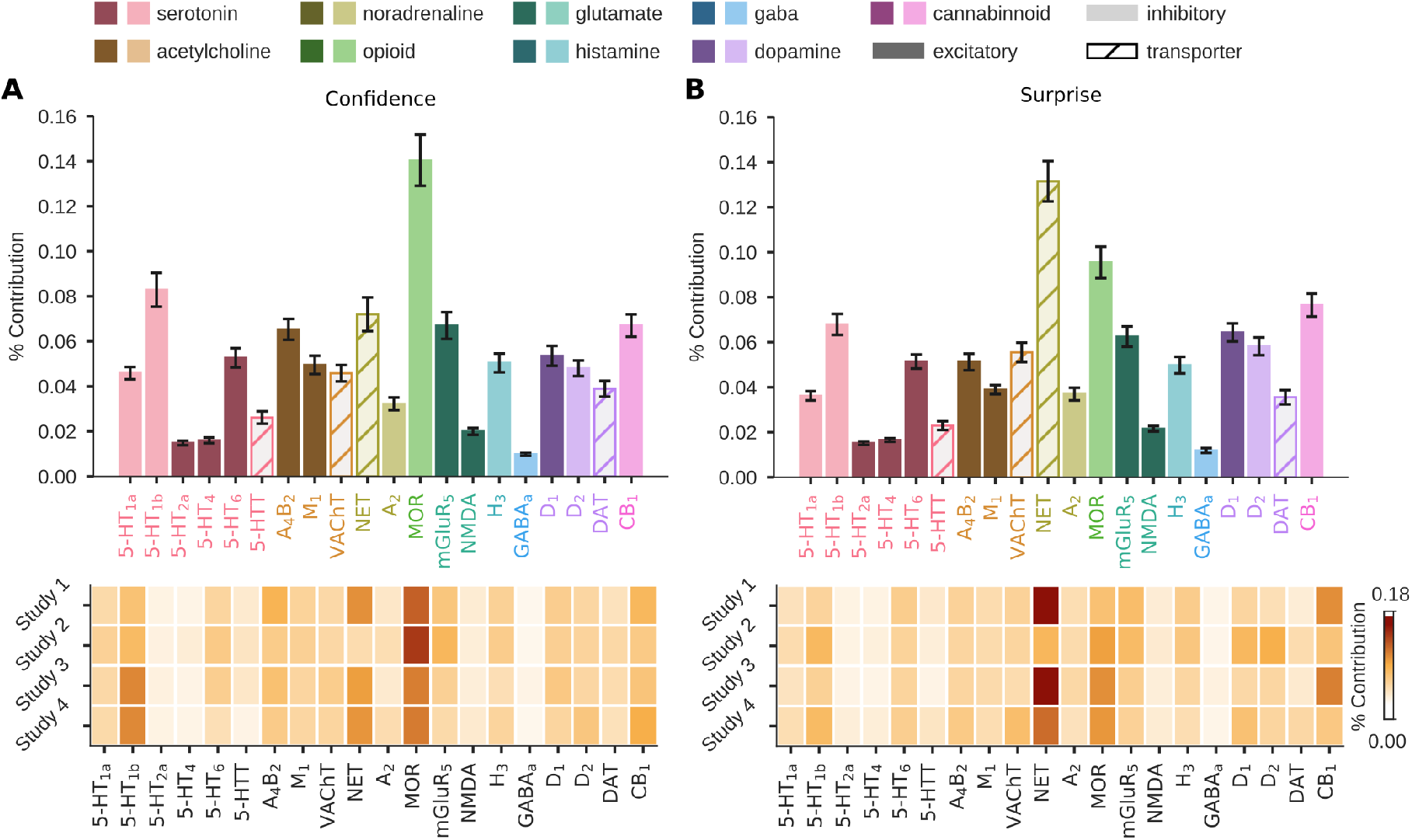
Contribution of each receptor/transporter to explaining the learning-related effect maps. Dominance analysis assigns a proportion of the model fit to each independent variable. For comparison across studies and learning-related variables, each contribution is normalized by the total model fit, and expressed as a percentage. The top row in A) (confidence) and B) (surprise) depicts the mean percent contribution across all subjects and studies. Error bars reflect the SEM across all subjects. The heat maps illustrate the mean dominance results by study.

For surprise effect maps, the topography of the norepinephrine transporter (NET) made the largest contribution to the model fit (Figure 4B). Its contribution was weaker in Study 2, likely due to partial brain coverage and modality-specific activations in temporal regions. In the full regression model, the NET coefficient was positive and significant across subjects in each probability learning study (studies 1-3; FDR-corrected p <0.05), indicating that for probability learning regions with higher NET density were more activated by higher surprise, when taking all other receptors/transporters into account (Table 4). Several other receptors (5-HT_6_, D_1_, D_2_, CB_1_) also showed significant and consistent regression weights for each probability learning study but made smaller contributions to model fit. In the reward learning study, the NET effect appeared to be reversed compared to the probability learning studies, meaning that regions with lower NET density show larger surprise effects (Table 4). This reversal is compatible with the fact that surprise maps in the probability and reward tasks are generally uncorrelated.

Overall, these findings show that consistent receptor-level contributions can be identified across studies, demonstrating that learning-related activity is systematically shaped by the brain’s underlying neuromodulatory architecture.

## 3 Discussion

Across four probabilistic learning studies, we found that the cortical topography of confidence was strongly invariant, and that surprise showed partial invariance. We hypothesized that such stable functional patterns arise, in part, from the brain’s underlying receptor architecture. Supporting this view, both confidence- and surprise-related maps were partly predicted by cortical receptor and transporter distributions, allowing us to identify specific neuromodulatory pathways that appear to contribute to these representations.

The strong invariance of confidence representations across all probability and reward learning tasks suggests engagement of domain-general neural mechanisms (for a similar question in metacognition, see: 38; 39). This is particularly striking when taking into account that confidence (or its inverse uncertainty) is known to drive learning rate in probability learning, whereas in magnitude learning, learning rate is modulated more strongly by change-point probability (40). Replication in additional magnitude-learning paradigms will be important, but the current evidence points to shared neural mechanisms for confidence across tasks with distinct computational demands.

Adaptive learning is not shaped solely by confidence and surprise. Other computational variables such as environmental volatility (e.g., 25; 40) or predictability (21) also shape the updating of the learned estimate. However, these variables typically do not yield the stable and large-scale cortical patterns required for the present approach. More generally, the present results illustrate that when a computational variable expresses a domain general topography in the cortex, it becomes possible to probe the neuromodulatory basis of this topography.

Specifically, we interpret the invariance of both confidence and surprise maps partly as a consequence of constraints imposed by common neuromodulatory processes. Our findings reinforce growing evidence that receptor architecture shapes brain function (14; 17; 18; 19). However, our work specifically anticipated such a relationship given longstanding proposals that neuromodulatory projections are central to learning under uncertainty (7; 8; 9). Our framework allowed us to probe the potential specificity of this receptor–function relationship. The associations between receptor architecture and functional topography were substantially weaker for the language network, which is not thought to rely strongly on neuromodulatory systems. This demonstrates that the methodological approach is not limited to learning, but provides a general framework for assessing the extent of neuromodulatory involvement across cognitive domains and for identifying specific receptor and transporters that may shape a given process.

We found that the topography of specific receptors consistently predicted the topography of learning-related effects. Confidence was strongly associated with the *µ*-opioid receptor (MOR). Although MOR activation has diverse downstream effects, its inhibition of the locus coeruleus via endogenous opioids (41; 42) and its indirect disinhibition of dopamine neurons (43; 44) may be most relevant in the context of learning. The novel association between confidence in learning and cortical MOR has not been established previously and opens a promising avenue for future research. The opioid system has also been implicated in cognitive control (45; 46), suggesting a broader opioidergic role in cognition. The overall alignment between confidence effect maps and receptor densities was preserved even in the reward-learning task, suggesting that the invariant topography of the effect of confidence may, in part, reflect a domain-general neuromodulatory process.

The topography for surprise in probability learning was strongly associated with the norepinephrine transporter (NET). NET regulates the reuptake of extracellular norepinephrine, a neuromodulator proposed to modulate neural gain (47). Its importance in probability learning suggested by our results is consistent with influential theories on the involvement of norepinephrine in probabilistic learning (9) and previous pharmacological studies (48; 49). Interestingly, specific polymorphisms of the NET gene have been reported to account for individual differences in learning rate (50). Together, these results point to norepinephrine and its regulation via NET as being key to adaptive learning.

Beyond these systems, other neuromodulators revealed weaker but consistent associations across datasets. Specifically, serotonergic receptors (confidence and surprise), dopaminergic receptors (surprise), and acetylcholinergic receptors (confidence) accounted for the topography of learning-related effects, providing convergent support for neuromodulatory theories of learning under uncertainty (7; 8; 49).

Surprise effects showed a different topography, and a different association with receptors and transporters in the reward-learning task compared with the probability-learning tasks. The two maps overlapped in a core region of the right precentral sulcus and inferior frontal gyrus pars opercularis, but otherwise differed markedly. This difference requires replication with other reward-learning paradigms, but several task features likely contribute. First, in reward learning the sign of the prediction error has a strong affective component, as it denotes outcomes that are better or worse than expected. By contrast, prediction errors in the probability-learning tasks were affectively neutral. Second, the reward-learning task combined learning with decision-making, whereas the probability-learning tasks involved pure learning. Finally, in the reward-learning data, confidence and surprise were more strongly correlated across trials than in the probability-learning tasks, which may account for the stronger correlations of the surprise and confidence maps observed in the reward learning task.

The present findings should be interpreted in light of several limitations. First, our conclusions are constrained by limitations inherent to the PET receptor density data. We restricted our analyses to the cortex because PET radioligand sensitivity differs markedly between cortical and subcortical structures (14; 51; 52). Moreover, although the PET dataset we used (14; 32) represents the most comprehensive collection of receptor densities to date, it includes only a subset of all known receptors and transporters. The receptor maps were derived from different individuals than those contributing to the fMRI data. Future studies combining subject-specific PET and fMRI measurements could more directly test receptor–function relationship specifically for the *µ*-opioid receptor or the norepinephrine transporter. However, note that such studies would necessarily be limited in scope, as collecting multiple PET scans per participant is costly, logistically challenging, and ethically difficult. Together, this would make it impractical to survey the broad receptor and transporter space examined here. Another limitation of PET is its relatively low spatial resolution, which produces smooth maps that inflate apparent relationships and constrain analyses to a whole-brain scale. The development of higher resolution maps in the future might provide complementary insights into spatial dynamics.

Second, the linear model only explains a modest fraction of the explainable variance. Beyond suggesting the existence of other constrains (20), this most likely results from our rigorous approach: variance estimates were obtained from cross-validated, subject-level voxelwise analyses. Model performance would increase when using (smoother) group-level analyses, as it has been done in previous studies investigating a relationship between receptor distributions and brain function (14; 17; 18). Further, our analyses assumed linear effects of receptor and transporter densities, a conservative simplification that ignores the nonlinear and interactive nature of neuromodulatory signaling. To address this, we repeated the analysis including quadratic terms for all receptors and transporters. The results remained qualitatively unchanged and supported the same main conclusions (Figure 7. Models including interaction terms explain more (cross-validated) variance (see table 2). Future work should therefore investigate higher-order interactions among receptors and neurotransmitters, for which the current findings provide an empirical foundation.

Finally, the study is correlational in nature. Specifically, our analysis targets receptors and transporters whose spatial distributions constrain the cortical topography of learning-related effects; other receptors or transporters (particularly those with relatively homogeneous cortical distributions) may still play a role without imposing such spatial constraints. Nonetheless, our results complement causal evidence from pharmacological manipulations (48; 49) and extend the pharmacological approach by jointly evaluating multiple neuromodulatory systems. In doing so, we not only reinforce neuromodulatory accounts of adaptive learning but also identify new specific receptor-level hypotheses that can be tested in future experimental and pharmacological studies.

In summary, we showed that distinct receptor and transporter patterns underlie the topography of learning-related activity in the brain. Our results demonstrate that linking fMRI effect maps to receptor/transporter topographies makes it possible to identify neuromodulatory pathways associated with specific computational variables, such as confidence and surprise. These findings provide a framework for probing the neuromodulatory basis of cognition and for developing receptor/transporter-level theories of cognitive function.

## 4 Methods and Materials

All code that was used to perform the analyses can be found at https://github.com/TheComputationalBrain/NeuroMod.

### 4.1 fMRI datasets

To the best of our knowledge, subject-level fMRI effects maps for surprise and confidence in learning tasks (estimated while controlling for other factors, see below) are not openly available. We therefore first computed subject-level effect maps associated with confidence and surprise from a few datasets from which we could obtain the raw data in order to ensure consistent analysis methods across studies. All datasets stemmed from experiments that were approved by an Ethics Committee (Study 1: CPP-100032 Ile-de-France VII, Study 2: CPP-100055 Sud-Ouest et Outre Mer III; Study 3: CPP 08–021 Ile de France VII; Study 4: CPP-100-050 Ile de France III), and subjects gave informed consent prior to participating.

Specifically, we analyzed 4 fMRI datasets of learning under uncertainty that differed in terms of sensory modality (visual vs auditory), structure (Bernoulli vs. Gaussian distribution vs. transition probabilities), and estimate to learn (probability vs magnitude). Studies 1-3 were probability learning tasks (Figure 1A), and Study 4 was a learning and decision-making task (Figure 1B). In all tasks, the latent variable to learn could change suddenly and unpredictably and subjects were informed about the generative process underlying the sequence in a non-technical way.

#### 4.1.1 Study 1

comprises 7T fMRI data from (21). 26 participants were presented with a visual sequence composed of two Gabor patches without perceptual ambiguity drawn from a Bernoulli process. Occasionally, participants reported their estimate for one of the stimuli as well as their confidence about this estimate.

#### 4.1.2 Study 2

consists of yet unpublished 3T data in which 55 participants were presented with an auditory sequence of two tones. The tones were determined by a Bernoulli process and participants were occasionally asked to indicate their estimate for the probability of a sound and their confidence in this estimate. This study did not provide full brain coverage, as it focused on subcortical structures (mean coverage: 74.2%, SEM: 0.6). Six subjects were removed for measures of explained variance, due to significantly lower coverage than the rest of the subjects (mean coverage: 54.2%, SEM: 4.8).

#### 4.1.3 Study 3

was previously published in (22) and consists of 3T data from 21 participants learning the transition probabilities between two stimuli in the presented sequence. Both auditory and visual sequences were used in distinct sessions. Participants occasionally reported their confidence in the next stimulus given the identity of the previous one.

#### 4.1.4 Study 4

consists of yet unpublished 3T data from a magnitude (reward) learning task described in (23). 56 participants performed a sequence of choices in a two-armed bandit task trying to maximize cumulative reward, which required learning the underlying reward level for two choices with independently evolving reward distributions. They were presented with a reward between 1 and 100 points for the chosen option. Subjects were asked occasionally to make explicit reports of their reward level estimate as well as confidence about this estimate.

### 4.2 Processing and analysis

For each dataset, functional data were normalized to MNI space (MNI-ICBM 152 nonlinear 2009 template). Voxel time series were detrended, high-pass filtered (1/128 Hz), z-scored per experimental run, and spatially smoothed with a 5 mm FWHM Gaussian kernel. Analyzing raw rather than group-level data allowed us to apply identical regression models across all studies with the aim to derive effect maps for confidence and surprise based on an ideal observer model.

Effect maps correspond to the regression coefficients from a general linear model (GLM) obtained with Nilearn v0.11.1 (53). For all studies, the GLM modeled stimulus onset with confidence and surprise as trial-wise parametric modulators derived from the ideal Bayesian observer. It also included the questions asked in the task as well as several corresponding parametric modulations (reaction times, the ideal observer’s prediction and confidence as well as the subject’s prediction and confidence in response to the question, if applicable) and motion parameters.

In addition, we included other covariates to make the estimation of the effects of surprise and confidence more specific: the probability learning tasks included the model-derived current prediction of the upcoming stimulus *p* and predictability (corresponding to entropy: − *plog*_2_ (*p*) − (1 − *p*)*log*_2_(1 − *p*)), while the reward learning task included the expected reward difference between chosen and unchosen options, as estimated by the ideal observer.

### 4.3 Bayesian model of learning

Learning-related variables were derived from an ideal observer model, providing a principled and consistent reference across subjects and studies. The ideal observer model was implemented with a well-known solution based on a Hidden Markov Model (2; 54). In short, the model provides optimal estimates of the changing quantity of interest (item probability or reward level) *θ*_*t*_ given the observations so far, *y*_1:*t*_. This is achieved by iteratively combining the likelihood of the current observation and a prior:

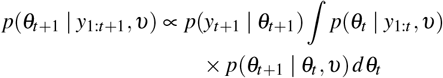

The learner assumes that *θ* can change unexpectedly with some prior probability *υ*, as captured by *p*(*θ*_*t*+1_ *θ*_*t*_, *υ*), in which case it is resampled uniformly. The volatility *υ*, the transition matrix and prior probability or reward levels (when applicable) correspond to the generative structure of the task. The prediction of the next stimulus is the average of the posterior distribution:

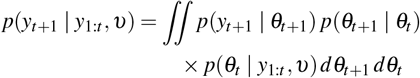

We formalized the confidence about the prediction for the next stimulus as the log precision of the distribution in the probability learning task

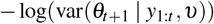

and as the maximum a posteriori probability in the decision task (this difference does not matter in practice). We quantified surprise as log improbability of the observation (28):

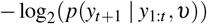

### 4.4 Null models

To assess the statistical significance of the correlations of fMRI effect maps, and of the prediction accuracy of fMRI effect maps from PET receptor/transporter density maps, we employed a spatial permutation approach known as the ‘spin test’. In this procedure, volumetric cortical maps were projected onto the spherical surface of an inflated brain (fsaverage6 template, 40,962 vertices per hemisphere) and randomly rotated. The rotated map replaced the original empirical map in the analysis, and this step was repeated 1000 times to generate a null distributions (for details, see 29; 30; 31). This test provides a stringent null model because the rotated maps preserve both the spatial autocorrelation and the value distribution (histogram) of the empirical data.

### 4.5 Overlap of effect maps

To assess the invariance of the effect maps for confidence and surprise, we combined individual participant regression coefficients in a group-level one-sample *t*-test against zero for each study separately. The resulting statistical maps were thresholded at *p* <0.001 (voxelwise) and corrected for multiple comparisons across voxels at the cluster level using family-wise error correction (*p*_FDR_ <0.05) based on Monte Carlo simulations implemented in Nilearn (53). To illustrate spatial invariance, we visualized the spatial overlap of significant positive and negative clusters across tasks separately (Figure 3B,E).

To complement this threshold-based approach with a quantitative, threshold-free metric, we computed Pearson correlations between unthresholded group-level effect maps of each study for both confidence and surprise (Figure 3C,F). To determine whether correlations between studies exceeded chance levels, we compared the observed correlations to a null distribution generated with the spatial permutation (‘spin test’) procedure. Multiple comparisons across different study-pairs were controlled using false discovery rate (FDR) correction.

### 4.6 PET receptor density maps

As a measure of cortical receptor distribution, we combined PET-derived whole-brain density maps for 19 unique neurotransmitter receptors and transporters recently made publicly available by Hansen et al. (14) with an additional *α*_2_-adrenergic receptor density map provided by Laurencin et al. (32). The resulting dataset spanned nine neurotransmitter systems. All PET images were available in voxel space, registered to the MNI-ICBM 152 nonlinear 2009 template, and averaged across participants within each study. When multiple PET maps for the same receptor were available (i.e., studies using the same tracer), they were combined using a weighted average. Each resulting receptor or transporter density map was then z-scored. Pairwise correlations among receptor maps are shown in Figure 6. For detailed acquisition and processing procedures, see the original publications (14; 32).

### 4.7 Relating brain activity and receptor density maps

We examined the relationship between the cortical distribution of neurotransmitter receptors and transporters and the fMRI effects associated with learning-related variables. Because our null model relies on surface-based rotations, all measures of explained variance were computed from cortical surface projections to ensure comparability.

#### 4.7.1 Average explained variance

To estimate the average explained variance within each study, we used a leave-one-subject-out cross-validation approach. For each study and variable, the model was trained on all but one participant and used to predict the held-out participant’s effect map. Model performance *R*^2^ was compared against the mean of a subject-level null distribution generated by repeating the same cross-validation procedure with spatially permuted receptor/transporter maps (1000 random rotations; ‘spin test’). Significance was assessed using paired *t*-tests between true and mean null *R*^2^ values across subjects, followed by false discovery rate (FDR) correction for multiple comparisons across studies. No null distribution was computed for Study 2 due to incomplete cortical coverage, which would have rendered the spatial permutations inconsistent with the original predictors.

#### 4.7.2 Maximum explainable variance

To contextualize model performance, we also estimated the maximum explainable variance for each study and variable. This was obtained by calculating the mean *R*^2^ of models that predicted each participant’s fMRI effect map from the group-mean map of all other participants within the same study and variable combination.

#### 4.7.3 Language network as a comparison

To establish a reference for comparison, we replicated our analysis on a dataset representing a cognitive function without a specific neuromodulatory hypothesis. We selected the language network because it meets this criterion and is characterized by robust, brain-scale, and highly reproducible activation patterns across individuals (similar to the effect maps for confidence and surprise). Specifically, we used publicly available subject-level contrast maps from 806 adults for a language *>* control contrast (for details, see 33)). To match the sample size of the largest learning dataset, we randomly selected 60 participants and applied the same analysis pipeline used for the learning studies. To test the robustness of the conclusions, we repeated the analysis with a different random subset of participants, which yielded the same conclusions.

#### 4.7.4 Explained variance ratio

We quantified the relative contribution of receptor and transporter densities by expressing the average explained variance as a proportion of the maximum explainable variance, which we refer to as the explained variance ratio. We conducted our primary analyses at the group level because several participants exhibited negative values for both explained and maximum explainable variance, rendering subject-level ratios difficult to interpret. To evaluate whether the explained variance ratio differed between each learning-related latent variable and the language network, we performed a group-level non-parametric permutation test. Specifically, for each comparison we computed the empirical difference in group means (mean learning study ratio - mean language network ratio) and generated a null distribution by randomly permuting condition labels across subjects (10,000 permutations). The one-sided p-value was obtained as the proportion of permuted differences that exceeded the empirical difference, and values were corrected for multiple comparisons using FDR.

For completeness, we additionally report subject-level analyses in the supplementary table 3. Here we computed a ratio for each subject (explained / maximum explainable variance) and tested differences between learning variables and the language network using a Mann–Whitney U test (one-sided; p-value was corrected for multiple comparison using FDR), providing a subject-level confirmation of the group-level findings.

### 4.8 Identifying receptors of interest

To identify the receptors or transporters that contributed most to the fit between receptor/transporter topographies and cortical fMRI effect maps, we applied dominance analysis (37) at the subject level (across voxels in volume MNI space), a method previously employed in similar studies (14; 55; 56). Dominance provides an interpretable measure of the contribution of each regressor by partitioning the model fit among predictors while accounting for dependence or collinearity between predictors. Specifically, the regression model is fit for every possible combination and subset of predictors (2^*p*−1^ regression models for *p* independent variables, here *p* = 20). The contribution of a given receptor/transporter is computed as the average increase in model fit across all models that include that receptor/transporter compared to models without it. The resulting values were then normalized by the full model *R*^2^ (i.e., the model including all 20 receptors and transporters) to allow comparison across latent variables (confidence and surprise) and studies (see also 14). The resulting values are reported as receptor/transporter contribution.

We also performed a variant of the analysis that included both linear and quadratic terms for each receptor/transporter (Figure 7). In this case, the contribution of each receptor/transporter was evaluated by grouping linear and quadratic terms, while all other aspects of the procedure remained the same.

Because dominance analysis does not account for the direction (sign) of the effects of predictors, we additionally examined regression weights from the full model including all 20 receptors and transporters. Significance of each predictor was assessed using a one-sample t-test against zero across subjects within each study. We report significant parameter estimates that exhibited a consistent sign across studies (*p*_FDR_ <0.05) in Table 4. These regression weights serve as supporting information, allowing interpretation of the direction of receptor contributions identified by the dominance analysis.

## 5 Acknowledgements

This work was supported by funding from the European Research Council (ERC grant 947105) to F.M. *α*_2_ receptor density map © Copyright CRNL, CERMEP – Imagerie du vivant, www.cermep.fr and Hospices Civils de Lyon. All rights reserved.

## Appendix

**Table 1.** Mean results by study for a linear regression analysis comparing explicit subject reports and ideal observer estimates.

| | | $R^2$ | coef | sd | $p$ |
| --- | --- | --- | --- | --- | --- |
| Study 1 | confidence | 0.038 | 0.105 | 0.109 | $4.70 \times 10^{-05}$ |
| | prediction | 0.641 | 0.961 | 0.291 | $3.84 \times 10^{-15}$ |
| Study 2 | confidence | 0.053 | 0.136 | 0.121 | $7.83 \times 10^{-11}$ |
| | prediction | 0.563 | 0.724 | 0.324 | $4.53 \times 10^{-22}$ |
| Study 3 | confidence | 0.059 | 0.177 | 0.120 | $1.44 \times 10^{-06}$ |
|  | prediction | not applicable (see text) |  |  |  |
| Study 4 | confidence | 0.109 | 0.425 | 0.236 | $4.65 \times 10^{-19}$ |
|  | prediction | different comparison (see text) |  |  |  |

Note that probability is expressed on the same scale in the ideal observer and subjects, making the regression coefficient interpretable and equal to 1 had subjects been ideal. By contrast, confidence is expressed in precision unit for study 1, 2 and 3, whose scale is in theory not bounded, whereas subject reported their confidence on a qualitative scale that was arbitrarily remapped to the unit interval for the purpose of the analysis; therefore the corresponding regression coefficient is not interpretable (and much lower than 1 due to the remapping).

**Figure 5.**
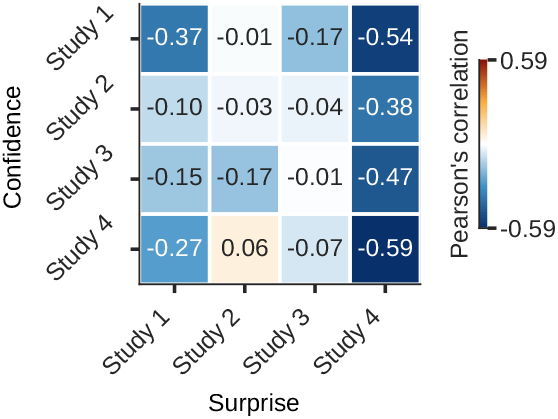
Correlations between confidence and surprise maps.

**Table 2.** Comparison of average model fit for different model complexities.

|  |  | linear | linear +<br>quadratic | linear +<br>interaction | full polynomial<br>model (2) |
| --- | --- | --- | --- | --- | --- |
| Confidence | Study 1 | 0.036 | 0.044 | 0.066 | 0.067 |
|  | Study 2 | 0.023 | 0.027 | 0.040 | 0.042 |
|  | Study 3 | 0.036 | 0.048 | 0.072 | 0.074 |
|  | Study 4 | 0.009 | 0.010 | 0.016 | 0.017 |
| Surprise | Study 1 | 0.047 | 0.057 | 0.122 | 0.127 |
|  | Study 2 | 0.036 | 0.046 | 0.102 | 0.105 |
|  | Study 3 | 0.022 | 0.027 | 0.048 | 0.050 |
|  | Study 4 | 0.018 | 0.020 | 0.040 | 0.041 |

All $R^2$ reflect cross validated results

**Table 3.** Results from Mann–Whitney U test on subject level explained variance ratios.

|  |  | U | p (FDR corrected) |
| --- | --- | --- | --- |
| Confidence | Study 1 | 1171.0 | 0.0006 |
|  | Study 2 | 2036.0 | 0.0006 |
|  | Study 3 | 986.0 | 0.0006 |
|  | Study 4 | 2229.0 | 0.0169 |
| Surprise | Study 1 | 1008.0 | 0.0267 |
|  | Study 2 | 1831.0 | 0.0169 |
|  | Study 3 | 780.0 | 0.0699 |
|  | Study 4 | 2376.0 | 0.0025 |

Mann-Whitney U test on subject-level explained variance ratios (explained variance/maximum explainable variance) comparing each study/variable combination to the subject-level ratios from the language network data. Significance tests are one-sided and FDR corrected for multiple comparisons.

**Table 4.** Parameter estimates across receptors/transporters.

| Receptor | Confidence |  |  |  | Surprise |  |  |  |
| --- | --- | --- | --- | --- | --- | --- | --- | --- |
|  | Study 1 | Study 2 | Study 3 | Study 4 | Study 1 | Study 2 | Study 3 | Study 4 |
| 5HT1a | 0.0016 | 0.0022 | 0.0027 | 0.0001 | 0.0045 | 0.0006 | 0.0020 | -0.0002 |
| 5HT1b | **-0.0045** | **-0.0020** | **-0.0059** | **-0.0020** | 0.0023 | 0.0023 | 0.0003 | 0.0016 |
| 5HT2a | 0.0007 | 0.0019 | -0.0014 | -0.0007 | 0.0040 | 0.0047 | 0.0012 | -0.0004 |
| 5HT4 | -0.0017 | -0.0023 | -0.0004 | 0.0002 | -0.0028 | -0.0046 | -0.0019 | 0.0016 |
| 5HT6 | 0.0036 | 0.0039 | 0.0045 | 0.0016 | **-0.0105** | **-0.0020** | **-0.0045** | 0.0006 |
| 5HTT | 0.0020 | 0.0009 | 0.0023 | 0.0006 | -0.0012 | 0.0005 | 0.0011 | -0.0013 |
| A4B2 | **-0.0059** | **-0.0027** | **-0.0049** | **-0.0011** | 0.0024 | -0.0020 | -0.0003 | 0.0015 |
| CB1 | **0.0073** | **0.0031** | **0.0061** | **0.0013** | **-0.0097** | **-0.0028** | **-0.0055** | **-0.0017** |
| D1 | 0.0027 | -0.0012 | 0.0034 | 0.0006 | **-0.0039** | **-0.0040** | **-0.0016** | -0.0004 |
| D2 | -0.0011 | -0.0013 | -0.0023 | -0.0006 | **0.0047** | **0.0057** | **0.0032** | 0.0001 |
| DAT | 0.0008 | 0.0015 | 0.0003 | 0.0007 | -0.0009 | 0.0003 | -0.0005 | 0.0004 |
| GABAA | -0.0004 | -0.0004 | -0.0001 | 0.0000 | -0.0005 | -0.0007 | -0.0005 | -0.0002 |
| H3 | 0.0017 | 0.0011 | 0.0018 | 0.0006 | -0.0013 | -0.0015 | 0.0000 | 0.0005 |
| M1 | -0.0020 | -0.0015 | -0.0024 | -0.0007 | 0.0006 | -0.0029 | 0.0001 | -0.0006 |
| mGluR5 | 0.0001 | 0.0015 | -0.0033 | 0.0006 | -0.0031 | 0.0053 | 0.0006 | 0.0010 |
| MOR | **-0.0070** | **-0.0051** | **-0.0066** | -0.0011 | 0.0028 | -0.0003 | 0.0006 | 0.0013 |
| NET | -0.0003 | -0.0013 | 0.0017 | -0.0001 | **0.0141** | **0.0043** | **0.0080** | -0.0019 |
| NMDA | -0.0002 | 0.0008 | 0.0015 | 0.0001 | 0.0009 | -0.0012 | -0.0001 | 0.0003 |
| VACHT | 0.0015 | 0.0020 | 0.0041 | 0.0010 | -0.0036 | 0.0001 | -0.0015 | -0.0016 |
| A2 | -0.0001 | -0.0008 | -0.0008 | -0.0003 | -0.0007 | -0.0007 | -0.0009 | -0.0008 |

Bold values indicate that regression weights are significant in the full regression model (FDR-corrected; $p < 0.5$ ) in at least all probability learning studies (studies 1-3) and have a consistent direction of effect across studies.

**Figure 6.**
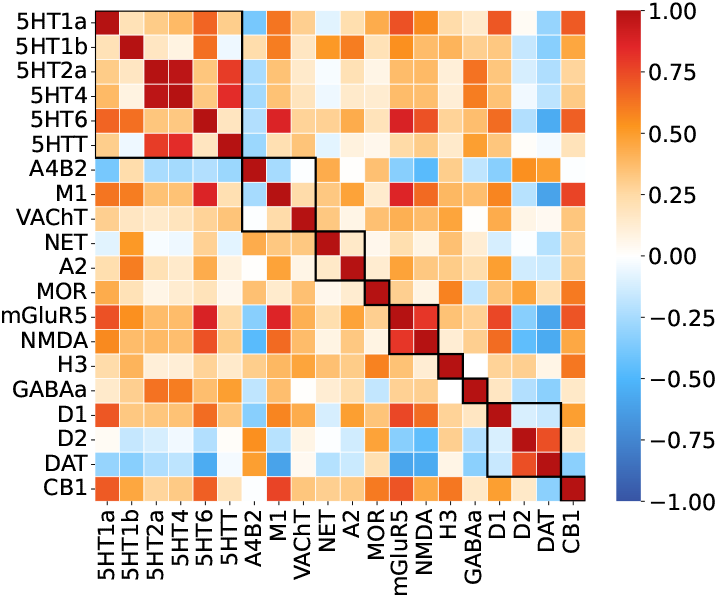
Pearson’s correlations between pairs of receptor/transporter distributions, grouped by neuromodulator.

**Figure 7.**
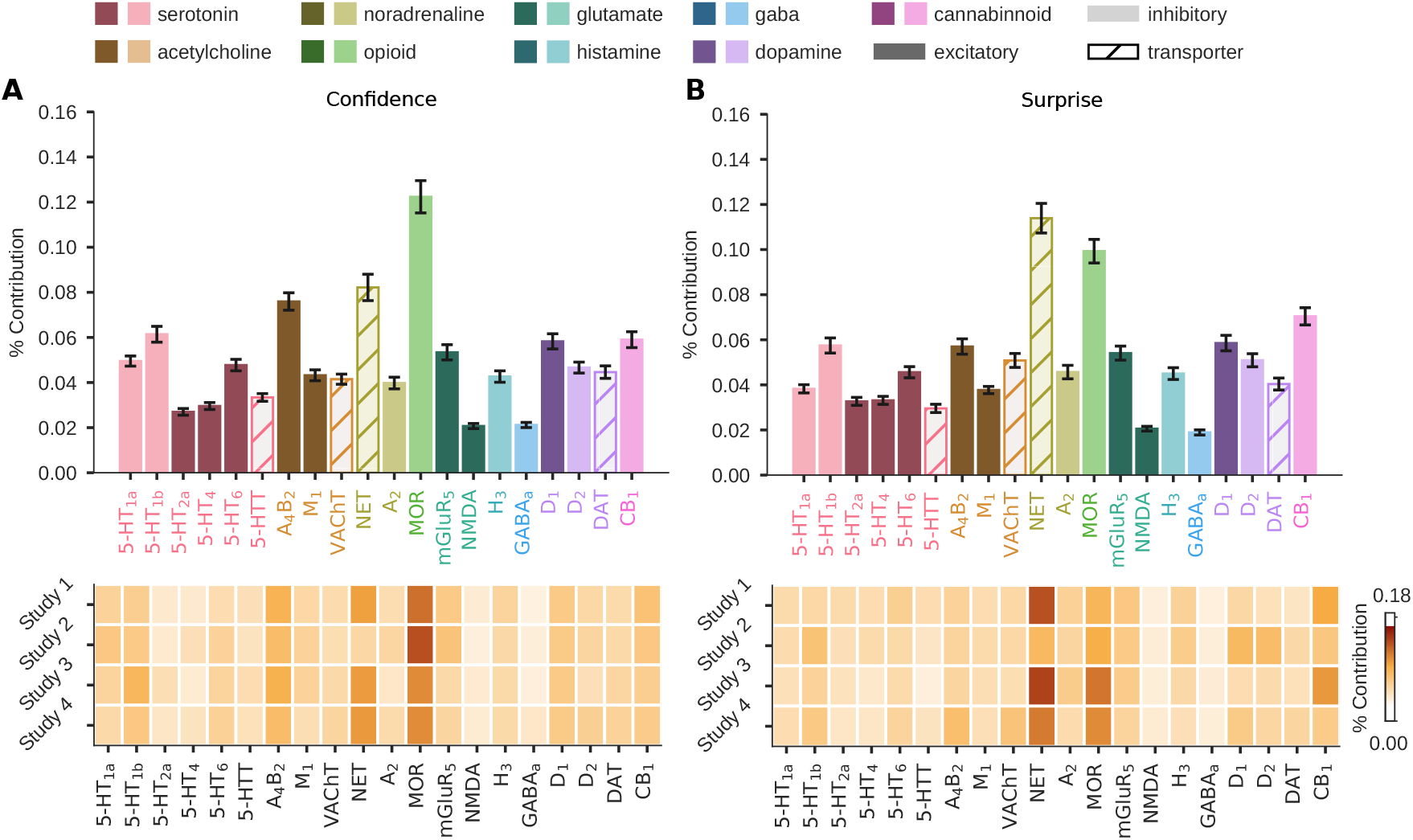
Results of the dominance analysis for a model including the linear and quadratic predictors, revealing the contribution of each receptor and transporter to the model fit. The percent contribution of each receptor reflects the variables dominance normalized by the model fit (*R*^2^). The top row in depicts the mean percent contribution across all subjects from all probability studies. Error bars reflect the SEM. The heat maps illustrate the mean dominance results by study.

## Notes

### Competing Interest Statement

The authors have declared no competing interest.

### Summary of Updates

Extended conference contribution to full manuscript.

